# Fatty acid bioconversion in harpacticoid copepods in a changing environment: a transcriptomic approach

**DOI:** 10.1101/797753

**Authors:** Jens Boyen, Patrick Fink, Christoph Mensens, Pascal I. Hablützel, Marleen De Troch

## Abstract

By 2100, global warming is predicted to significantly reduce the capacity of marine primary producers for long-chain polyunsaturated fatty acid (LC-PUFA) synthesis. Primary consumers such as harpacticoid copepods (Crustacea) might mitigate the resulting adverse effects on the food web by increased LC-PUFA bioconversion. Here, we present a high-quality *de novo* transcriptome assembly of the copepod *Platychelipus littoralis*, exposed to changes in both temperature (+3°C) and dietary LC-PUFA availability. Using this transcriptome, we detected multiple transcripts putatively encoding for LC-PUFA-bioconverting front-end fatty acid desaturases and elongases, and performed phylogenetic analyses to identify their relationship with sequences of other (crustacean) taxa. While temperature affected the absolute fatty acid concentrations in copepods, LC-PUFA levels remained unaltered even when copepods were fed a LC-PUFA-deficient diet. While this suggests plasticity of LC-PUFA bioconversion within *P. littoralis*, none of the putative front-end desaturase or elongase transcripts were differentially expressed under the applied treatments. Nevertheless, the transcriptome presented here provides a sound basis for future ecophysiological research on harpacticoid copepods.

## Introduction

Global climate change and the resulting increase in sea surface temperature over the past decades have profoundly impacted marine organisms and ecosystems (1). This trend is likely to continue for the next decades, with a projected global mean sea surface temperature (SST) increase of 2.73°C by 2090-2099 compared to 1990-1999 levels according to the business-as-usual Representative Concentration Pathway (RCP) 8.5 (2). This temperature rise affects the physiological performance and fitness of marine organisms and consequently triggers adverse changes in marine ecosystems as well as the goods and services they provide (3). Indeed, prominent climate-related shifts in nutrient and food supplies have already been observed in coastal areas worldwide (4,5). At the base of marine food webs, global warming is predicted to strongly impair the production of key nutritional fatty acids (FAs) by primary producers such as diatoms. These microalgae, like all living organisms, alter their FA composition due to temperature-dependent cell membrane restructuring, a process known as homeoviscous adaptation (6,7). Specifically, proportions of long-chain polyunsaturated FAs (LC-PUFAs) such as eicosapentaenoic acid (20:5*ω*3, EPA) and docosahexaenoic acid (22:6*ω*3, DHA) in microalgae are projected to decline strongly in the coming century (7). While the impact varies between taxa and LC-PUFA compounds, overall a reduced LC-PUFA availability is expected for higher trophic levels, many of which strongly rely on dietary LC-PUFAs to fulfil their metabolic requirements (5,8). Given the important role of LC-PUFAs in structural and physiological processes and as precursors for hormones and signalling molecules (9), a reduced dietary LC-PUFA availability impacts growth, fecundity and fitness of consumers (10,11). The relative contributions of the direct (increased SST) and the indirect (decreased dietary LC-PUFA availability) effect of global warming on higher trophic levels remains yet understudied (12). Growth and FA composition of European sea bass *Dicentrarchus labrax* and European abalone *Haliotis tuberculata* were impacted by temperature but only to a lesser extent by diet (13,14). This lesser dependency on dietary LC-PUFAs was attributed to the endogenous bioconversion of short-chain saturated FAs into LC-PUFAs. Numerous animal species, many of which aquatic invertebrates, are known to have at least some capacity for LC-PUFA bioconversion to cope with dietary changes (15,16). Benthic harpacticoid copepods (Crustacea) are key primary consumers in (coastal) marine and estuarine sediments (17) and are known for their capacity for LC-PUFA bioconversion (18–21). Coastal and estuarine environments undergo strong and stochastic fluctuations in temperature and nutrient availability. Harpacticoid copepods already adapted to such environments might be able to cope with the effects of global warming due to their LC-PUFA bioconversion capacity (19). However, the environmental shifts driving this bioconversion are yet poorly understood (16). Insights in the molecular aspects of LC-PUFA bioconversion are therefore required to understand this pivotal toolbox in crustaceans.

Converting short-chain saturated FAs to LC-PUFAs is achieved by a series of desaturase and elongase enzymes, which introduce a double bond or add two C-atoms to the FA chain, respectively (16). Desaturase enzymes themselves can be split up in front-end and methyl-end (or *ω*-end) desaturases, depending on the location of the double bond insertion (22). For crustaceans, genes encoding for front-end desaturase and elongase enzymes were so far mainly detected in decapods (23–27). Interestingly, a recent study discovered genes encoding *ω*-end desaturases in many aquatic invertebrates including some orders of copepods, challenging the current dogma that *de novo* PUFA biosynthesis occurs exclusively in marine microbes (28). Nielsen et al. (2019) identified putative front-end desaturase genes in multiple copepod species (29). Understanding how changes in diet or temperature affect those genes at the transcriptomic (i.e. gene expression) level has so far only been investigated in cyclopoid copepods (29,30). Within the order of Harpacticoida, transcriptomic resources are so far only available for *Tisbe holothuriae* (BioProject PRJEB23629) and three species of the *Tigriopus* genus (31–33).

Given the ecological importance of benthic harpacticoid copepods at the plant-animal interface, there is a need to better characterize their physiological response to global change at the molecular level. To do so, we investigated the transcriptomic and FA-metabolic response of the benthic harpacticoid copepod *Platychelipus littoralis* (Brady, 1880) towards both direct and indirect effects of global warming within a multifactorial setting, combining a change in SST (current versus future scenario) with a change in the dietary LC-PUFA availability (LC-PUFA-rich diatoms versus LC-PUFA-deficient green algae as food sources (34,35)). *P. littoralis* is a common intertidal species in European estuaries, and temperature-dependent LC-PUFA turnover rates have been demonstrated previously using compound-specific stable isotope analysis, even within a short timeframe of six days (19). This study presents a high-quality *de novo* transcriptome assembly from *P. littoralis* and reveals differentially expressed (DE) genes and FA profile changes towards both diet and temperature. Furthermore, putative PUFA desaturase and elongase genes are identified and compared phylogenetically with genes of other crustacean species.

## Material and Methods

### 1. Experiment

*Nitzschia* sp. (strain DCG0421, Bacillariophyceae) and *Dunaliella tertiolecta* (Chlorophyceae) were obtained from the BCCM/DCG Diatoms Collection (hosted by the Laboratory for Protistology & Aquatic Ecology - Ghent University) and the Aquaculture lab - Ghent University, respectively. Both algae were non-axenically cultured at 15 ± 1°C in filtered (3 μm; Whatman Grade 6) and autoclaved natural seawater (FNSW), supplemented with Guillard’s (F/2) Marine Water Enrichment solution (Sigma-Aldrich, Overijse, Belgium) and NutriBloom Plus (Necton) for *Nitzschia* sp. and *D. tertiolecta*, respectively. Food pellets were prepared through centrifugation and lyophilisation and stored at −80°C. In parallel, quadruplicate algae samples were stored at −80°C for later FA analysis. Additional quadruplicate algae samples were filtered (Whatman GF/F) and lyophilized for particulate organic carbon determination using high temperature combustion. *Platychelipus littoralis* specimens were collected from the top sediment layer of the Paulina intertidal mudflat (Westerscheldt estuary, The Netherlands; 51°21’N, 3°43’E) in August 2018. After sediment sieving (250 μm) and decantation, live adults were randomly collected using a glass Pasteur pipette under a stereo microscope. Copepods were cleaned by transferring them thrice to Petri dishes with clean FNSW and were kept in clean FNSW overnight to allow gut clearance prior to the start of the experiment.

The ten-day experiment had a fully crossed design with the factors temperature (19 ± 1 or 22 ± 1°C) and diet (*Nitzschia* sp. or *D. tertiolecta*). Temperature levels were based on the current mean August sea surface temperature at the sampling location (data obtained from www.scheldemonitor.be) and a global sea surface temperature increase of 3°C by 2100 as predicted by RCP8.5 (36). The food sources were offered as pre-thawed, rehydrated food pellets at a concentration of 3.69 ± 0.22 mg C l^−1^ day^−1^ and 5.10 ± 1.16 mg C l^−1^ day^−1^ for *Nitzschia* sp. and *D. tertiolecta* respectively, which are considered non-limiting food conditions (19,37). The combinations of diet and temperature yielded four treatments, each consisting of Petri dishes (52 mm diameter) filled with ten ml FNSW incubated in TC-175 incubators (Lovibond), with each treatment having four replicates for transcriptomic analysis (100 copepods per Petri dish) and three replicates for FA analysis (50 copepods per Petri dish). Each day, copepods were transferred to new units containing new temperature-equilibrated FNSW and were offered new pre-thawed food pellets. Triplicate copepod samples (50 specimens each) were collected from the field similarly as the specimens used in the experiment, and were stored at - 80°C for analysis of the initial (field) FA composition. At the end of the experiment, mortality was assessed, and all live specimens were transferred to Petri dishes with clean FNSW to remove food particles from the cuticle. Copepods for transcriptomic analysis were immediately thereafter flash-frozen in liquid nitrogen and stored at −80°C. Copepods for FA analysis were stored overnight to allow gut clearance prior to storage at −80°C. Differences in copepod survival between diet and temperature treatments and due to initial density (100 vs. 50 copepods per Petri dish) was statistically assessed in R v.3.6.0 (38) using a type II three-way ANOVA, a Tukey normalization transformation, and a stepwise model selection by AIC.

### 2. FA analysis

FA methyl esters (FAMEs) were prepared from lyophilized algal and copepod samples using a direct transesterification procedure with 2.5 % (v:v) sulfuric acid in methanol as described by De Troch et al. (18). The internal standard (nonadecanoic acid, Sigma-Aldrich, 2.5 μg) was added prior to the procedure. FAMEs were extracted twice with hexane. FA composition analysis was carried out with a gas chromatograph (HP 7890B, Agilent Technologies, Diegem, Belgium) equipped with a flame ionization detector (FID) and connected to an Agilent 5977A Mass Selective Detector (Agilent Technologies, Diegem, Belgium). The GC was further equipped with a PTV injector (CIS-4, Gerstel, Mülheim an der Ruhr, Germany). A HP88 fused-silica capillary column (60m×0.25mm×0.20μm film thickness, Agilent Technologies) was used at a constant Helium flow rate (2 ml min^−1^). The injected sample (2 μl) was split equally between the MS and FID detectors using an Agilent capillary flow technology splitter. The oven temperature program was as follows: at the time of sample injection the column temperature was 50°C for 2 min, then gradually increased at 10°C min^−1^ to 150°C, followed by a second increase at 2°C min^−1^ to 230°C. The injector temperature was held at 30°C for 6 s and then ramped at 10°C s^−1^ to 250°C and held for ten min. The transfer line for the column was maintained at 250°C. The quadrupole and ion source temperatures were 150 and 230°C, respectively. Mass spectra were recorded at 70 eV ionization voltage over the mass range of 50-550 m/z units.

Data analysis was done with MassHunter Quantitative Analysis software (Agilent Technologies). The signal obtained with the FID detector was used to generate quantitative data of all compounds. Peaks were identified based on their retention times, compared with external standards as a reference (Supelco 37 Component FAME Mix, Sigma-Aldrich) and by the mass spectra obtained with the Mass Selective Detector. FAME quantification was based on the area of the internal standard and on the conversion of peak areas to the weight of the FA by a theoretical response factor for each FA (39,40). Statistical analyses were performed in R v.3.6.0 (38). Shapiro-Wilk test and Levene’s test were used to check for normal distribution and homoscedasticity. The non-parametric Wilcoxon rank sum test was used to test for the difference in absolute and relative concentration of the individual FA compounds between field and incubated copepods. The type II two-way ANOVA and the non-parametric Scheirer-Ray-Hare test were used to test for the effects of diet and temperature on the absolute and relative concentration for each FA compound. Multivariate statistics were performed to test the effects of incubation, diet and temperature on the overall FA composition. Non-metric multidimensional scaling (nMDS) and PERMANOVA were performed after square root transformation (Bray-Curtis dissimilarity). Mean values are presented with ± s.d. The FA shorthand notation A:BωX is used, where A represents the number of carbon atoms, B the number of double bonds, and X the position of the first double bond counting from the terminal methyl group.

### 3. Transcriptomic analysis

Total RNA from 45 to 97 pooled *P. littoralis* specimens per sample was isolated using the RNeasy Plus Micro Kit (QIAGEN) following an improved protocol (See Supplementary Methods). Total RNA was extracted from all four replicates from the two *Nitzschia* sp. treatments, but due to high copepod mortality, only three out of four replicates from each of the two *D. tertiolecta* treatments could be used. Total RNA quality and quantity were assessed by both a NanoDrop 2000 spectrophotometer (Thermo Scientific) and a 2100 Bioanalyzer (Agilent Technologies). cDNA libraries were constructed using the Illumina TruSeq Stranded mRNA kit and samples were run on an Illumina HiSeq 4000 platform with 75 bp paired-end reads at the Cologne Centre for Genomics (University of Cologne).

Read quality was assessed using FASTQC v.0.11.7. The reads were quality-trimmed and adapter-clipped using Trimmomatic (41), and the raw read files are available at the NCBI Short Read Archive under BioProject PRJNA575120. The transcriptome was assembled *de novo* using Trinity v.2.8.4 (42) including *in silico* read normalization. Multiple tools were used to assess assembly quality. First, the percentages of reads properly represented in the transcriptome assembly were calculated for each sample using Bowtie2 v.2.3.4 (43). Second, the transcripts were aligned against proteins from the Swiss-Prot database (February 16, 2019) using blastx (cutoff of 1e^−20^), and the number of unique proteins represented with full-length or nearly full length transcripts (>90 %) were determined. Third, we performed Benchmarking Universal Single-Copy Orthologs (BUSCO) v.3.1.0 analysis using the arthropod dataset to estimate transcriptome assembly and annotation completeness (44). Lastly, both the N50 statistic and the E90N50 statistic (using only the set of transcripts representing 90 % of the total expression data) were determined based on transcript abundance estimation using salmon v.0.12.0 following the quasi-mapping procedure (45). The transcripts were annotated using Trinotate (46), an open-source toolkit that compiles several analyses such as coding region prediction using TransDecoder (http://transdecoder.github.io), protein homology identification using BLAST and the Swiss-Prot database, protein domain identification using HMMER/Pfam (47,48), and gene annotation using EGGNOG, KEGG and Gene Ontology (GO) database resources (49–51). The transcriptome (GHXK00000000) is available at the NCBI TSA Database under BioProject PRJNA575120. Transcript abundance per sample was estimated using salmon v.0.12.0 following the quasi-mapping procedure (45). The R v.3.0.6 (38) package edgeR v.3.26.4 (52) was used to identify significantly DE transcripts. Transcripts with >5 counts per million in at least three samples were retained. TMM normalization was applied (53), and differential expression was determined using a gene-wise negative binomial generalized linear model with quasi-likelihood F-tests (glmQLFit) with diet, temperature and the diet x temperature interaction as factors. Transcripts with an expression fold change ≥2 at a false discovery rate ≤0.05 (Benjamini-Hochberg method (54)) were considered significantly DE. The R package topGO v.2.36.0 (55) was used to test for enriched GO terms for both the diet and the temperature treatment. The enrichment tests were run using the weight01 algorithm and the Fisher’s exact test statistic, with GO terms considered significantly enriched when p≤0.01.

A transcript was categorized as encoding a front-end desaturase when it contained the two essential Pfam domains *Cytb5* (PF00173) and *FA_desaturase* (PF00487), three diagnostic histidine boxes (HXXXH, HXXXHH, and QXXHH) and a heme-binding motif (HPGG) (22). A transcript was categorized as encoding an elongase when it contained the Pfam domain *ELO* (PF01151) and the diagnostic histidine box (HXXHH) (22). Nucleotide coding sequences of the transcripts were trimmed to those conserved regions and aligned using MAFFT v7.452 (56) with default parameters. Sequences that aligned badly, contained long indels or were identical to other sequences after trimming were removed. Additional well-annotated (sometimes functionally characterized) crustacean sequences found to be most closely related to each of the *P. littoralis* transcripts through blastx were added to the alignment. We also included desaturase and elongase sequences from the hydrozoan *Hydra vulgaris*, copepod front-end desaturase sequences from Nielsen et al. (2019) and human elongase sequences Elovl1 to Elovl7. For each gene family, an unrooted maximum likelihood phylogenetic tree was built using RAxML v.8.2.4 (57) with a General Time Reversible model of nucleotide substitution and CAT approximation. The final tree was rooted (midpoint), visualized and edited with FigTree v.1.4.3 (http://tree.bio.ed.ac.uk/software/figtree).

## Results

The two algal diets differed in their FA composition (Table S1). EPA and DHA were highly abundant in *Nitzschia* sp. (26.85 ± 2.26 % and 5.68 ± 0.29 %) while not detected in *D. tertiolecta*. On the contrary *D. tertiolecta* exhibited a high content of α-linolenic acid (ALA, 18:3*ω*3; 43.77 ± 1.29 %) compared to *Nitzschia* sp. (0.05 ± 0.03 %). Mean copepod survival after ten days was 75.46 ± 27.79 %. Besides high variation between replicates, no significant effects of diet, temperature, initial density or interactions on copepod survival were detected (three-way ANOVA; AIC = 1696.8; F_(7,20)_ = 1.382; p = 0.2665).

### 1. Changes in FA content and composition

At the end of the experiment, total FA content in copepods in the treatments (149.47 ± 21.37 ng copepod^−1^) declined compared to control values from the field (197.29 ± 4.50 ng copepod^−1^) (p = 4.4e^−3^). Overall FA composition of the field copepods was significantly distinct from the ones of the incubated copepods (F_(1,14)_ = 10.83; p = 3.0e^−3^) (Fig. 1). In the experiment, overall copepod FA composition was significantly affected by temperature (F_(1,11)_ = 4.30; p = 0.015) but not by diet (F_(1,11)_ = 1.83; p = 0.13, no interaction) (Fig. 1). Temperature but not diet significantly affected total FA content, which was lower at 22°C compared to 19°C (Fig. 2, Table S2). Temperature but not diet also significantly affected absolute concentrations of anteiso-15:0, 15:0, 16:0, 18:1*ω*7, 24:0, DHA and ∑MUFA, which were all lower at 22°C (Table S2). The diet only significantly affected the absolute ALA concentration, with copepods fed the ALA-rich diet *D. tertiolecta* having a higher absolute ALA concentration compared with copepods fed with *Nitzschia* sp. (Fig. 2, Table S2). A significant temperature x diet interaction was found for oleic acid (OA, 18:1*ω*9). Absolute OA concentration was higher at 19°C when fed *Nitzschia* sp., an effect that reversed (higher at 22°C) when fed *D. tertiolecta* (Fig. 2, Table S2). ∑PUFA, ∑SFA, or important LC-PUFAs such as EPA and ARA were affected by neither temperature nor diet. Despite the absence of EPA and DHA in *D. tertiolecta*, copepods fed this diet were able to maintain similar relative EPA and DHA concentrations as in copepods fed with *Nitzschia* sp. (Table S3).

**Figure 1.**
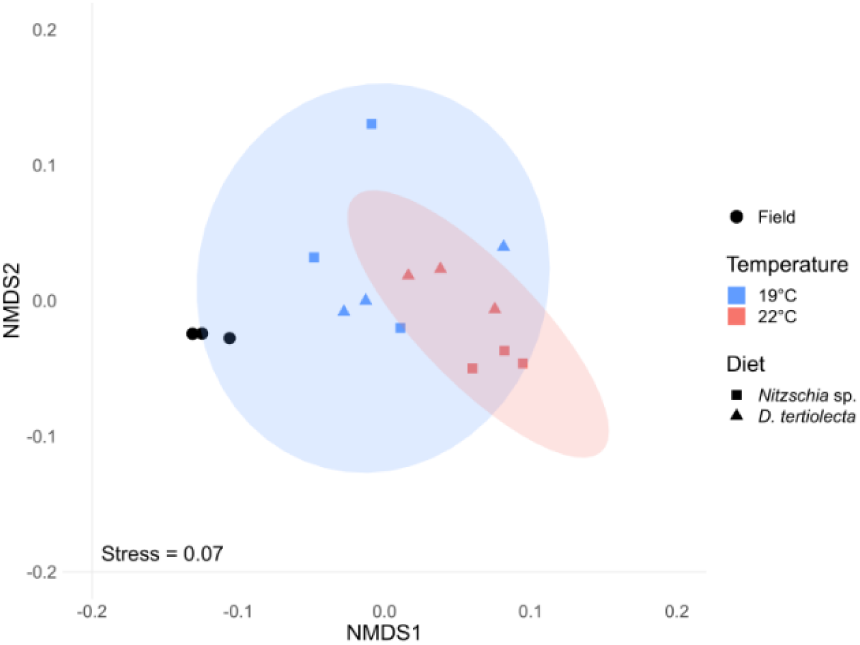
Non-metric multidimensional scaling (Bray-Curtis dissimilarity) on absolute FA concentrations (ng copepod^−1^) of the four treatments and field samples. Ellipses indicate 95 % confidence levels of the factor temperature.

**Figure 2.**
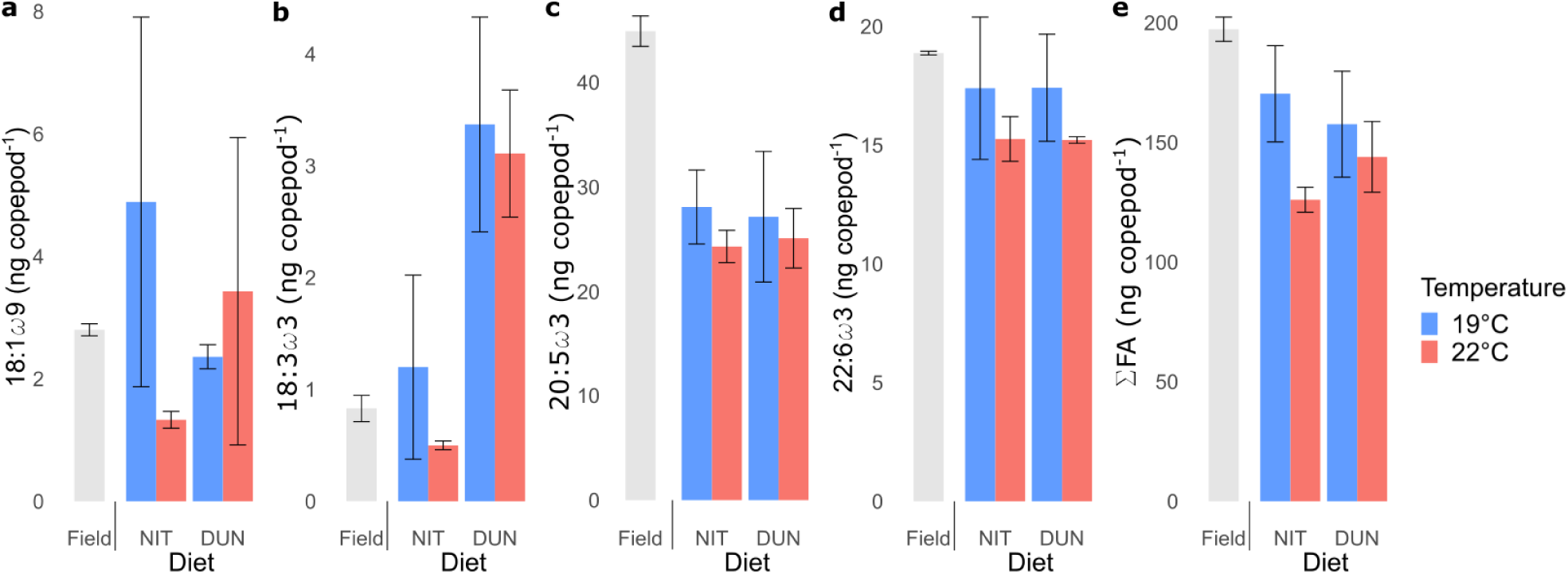
Mean absolute fatty acid concentration (ng copepod^-1^ ± s.d.; n=3) of *P. littoralis* prior (Field) and after ten days of incubation with *Nitzschia* sp. (NIT) or *D. tertiolecta* (DUN). **a** OA (18:1*ω*9) **b** ALA (18:3*ω*3) **c** EPA (20:6*ω*3) **d** DHA (22:6*ω*3) **e** ΣFA (total fatty acid concentration).

### 2. Transcriptome assembly and annotation

We sequenced 14 *P. littoralis* samples resulting in nearly 400 million paired-end Illumina reads. After quality filtering, an assembly was generated consisting of 287,753 transcript contigs (Table 1). 97.80 ± 0.44 % of the reads of each sample mapped back to the assembly, with 95.95 ± 0.68 % mapping as properly paired reads (aligning concordantly at least once). 7,088 proteins from the Swiss-Prot database were represented by transcripts with >90 % alignment coverage. 97.9 % of the BUSCO arthropod genes were represented by at least one complete copy (single: 26.1 %, duplicated: 71.8 %, fragmented: 1.1 %). The N50 and the E90N50 metrics were 1,360 and 2,657 respectively, and 90 % of the total expression data was represented by 35,540 transcripts. TransDecoder identified 296,142 putative open reading frame coding regions within the assembly, suggesting at least some transcripts contain multiple coding regions (Table 1). About a quarter of those putative ORFs were annotated with GO (24.9 %), KEGG (22.8 %) or EGNOGG terms (16.0 %) or were found to contain at least one Pfam protein family domain (29.5 %, Table 1).

**Table 1.** Assembly and annotation characteristics of the *P. littoralis* transcriptome

| Assembly and annotation characteristics | Value |
| --- | --- |
| Assembly |  |
| Number of raw reads | 395,794,835 |
| Average number of reads sample<sup>-1</sup> | 28,271,060 |
| Number of assembled bases (bp) | 207,643,686 |
| Number of assembled contigs | 287,753 |
| Average contig length (bp) | 721.6 |
| Median contig length (bp) | 343 |
| GC content (%) | 47.54 |
| Annotation |  |
| Number of ORFs | 296,142 |
| ORFs with GO terms | 73,836 |
| ORFs with KEGG terms | 67,484 |
| ORFs with EGNNOG terms | 47,358 |
| ORFs with Pfam protein family | 87,319 |

### 3. Differential expression and gene set enrichment analysis

After filtering out low expression transcripts, 24,202 transcripts were retained for DE analysis. Seven transcripts were DE between the two diet treatments (Fig. 3a,b, Table S4). Four of them were upregulated when copepods were fed with *Nitzschia* sp., while three were upregulated when copepods were fed with *D. tertiolecta*. 29 transcripts were DE between the two temperature treatments (Fig. 3c,d, Table S4). Five transcripts were upregulated at 19°C, while 24 were upregulated at 22°C. Four transcripts were DE in both treatments. All but two DE transcripts contained at least one putative ORF. Gene set enrichment analysis using topGO identified 22 and 30 enriched GO terms under the diet and temperature treatment, respectively (Table S5). Notable enriched functions in both treatments are related to microtubuli and cilia organization (Table S5). Certain terms detected in both treatments and primarily related to the biological process “glycerophospholipid biosynthetic process” and the molecular function “transferring acyl groups” (Table S5), were all attributed to the transcript *Plit_DN1805_c0_g1_i18*, which was downregulated when fed *D. tertiolecta* (compared to *Nitzschia* sp.) and upregulated at 22°C (compared to 19°C, Fig. 3, Table S4, S5). This transcript contains six ORFS which, according to blastx, match against the chicken protein acetoacetyl-CoA synthetase and the human protein glycerol-3-phosphate acyltransferase, both involved in the phospholipid metabolism.

**Figure 3.**
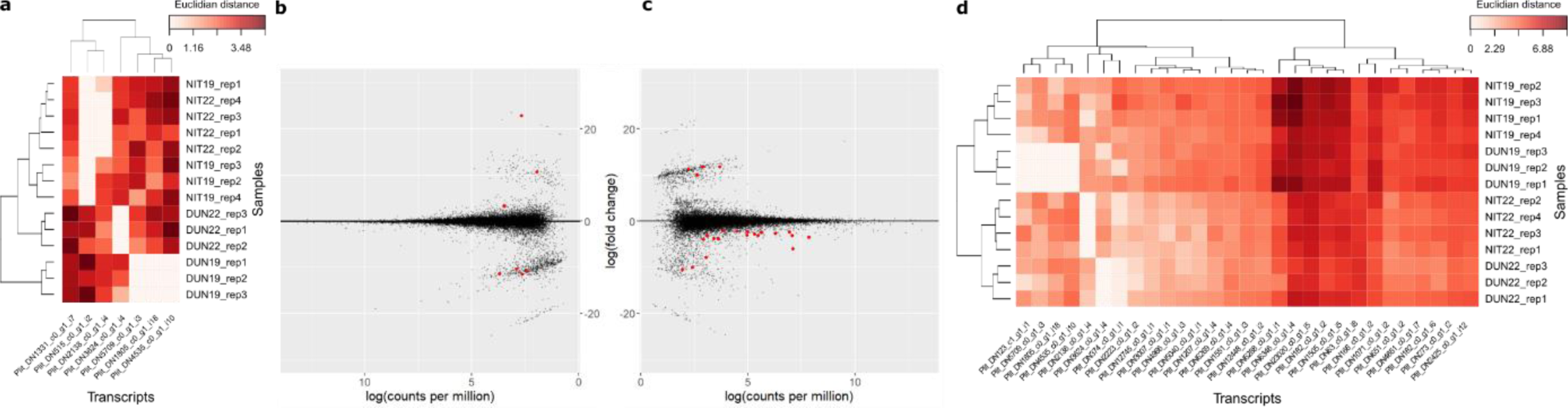
Heatmaps (**a**,**d**) and MA plots (**b**,**c**). MA plots show log fold change per transcript against its mean expression (in log counts per million) with diet (**b**) and temperature (**c**) as contrast. Red dots indicate significantly differentially expressed (DE) transcripts (i.e. log fold change ≥1 and false discovery rate ≤0.05). Black dots indicate non-significant DE transcripts. Heatmaps show relative expression level (Euclidean distance) of each significantly DE transcript in each sample with diet (**a**) and temperature (**d**) as contrast. Transcripts and samples are hierarchically clustered.

### 4. Identification and phylogenetic analysis of desaturase and elongase genes

Respectively 19 and 17 unique putative front-end desaturase and elongase sequences were identified in the *P. littoralis* transcriptome since they all exhibited diagnostic characteristics and aligned properly to other sequences. A maximum likelihood phylogenetic tree was built for each gene family (Fig. 4). Concerning the front-end desaturase sequences, we identified two distinct clades with high bootstrap support (Fig. 4a). The first clade contained five front-end desaturases from *P. littoralis* as well as from all other crustacean taxa. While one transcript is related to the functionally characterized Δ6 front-end desaturase sequence from the decapod *Macrobrachium nipponense*, the phylogenetic relationship of the other *P. littoralis* transcripts as a sister clade of the sequences of other copepod species is poorly supported (bootstrap value <50). The second clade contained only sequences from *P. littoralis* and one sequence of *H. vulgaris* in a basal position. The phylogenetic analysis of elongase transcripts produced multiple subclades (Fig. 4b). Two *P. littoralis* sequences and one *Daphnia magna* elongase sequence formed a clade with the human Elovl3 and Elovl6, classifying them as putatively Elovl6/Elovl3-like. One *P. littoralis* sequence, one *D. magnia* sequence and one *H. vulgaris* sequence were found to be closely related with the human Elovl4 and the Elovl4-like sequence from *Scylla paramamosain*, classifying them as putatively Elovl4-like. Four *P. littoralis* sequences and sequences from *D. magna, Caligus rogercresseyi* and *Lepeophtheirus salmonis* formed a subclade with the functionally characterized Elovl7 sequence from *Scylla olivacea* which is also a sister clade with the human Elovl7 and Elovl1, classifying those as putatively Elovl7/Elovl1-like. None of the *P. littoralis* sequences formed a clade with human Elovl5 and Elovl2, while ten *P. littoralis* sequences were not related with any of the sequences included in the analysis (Fig. 4b).

**Figure 4.**
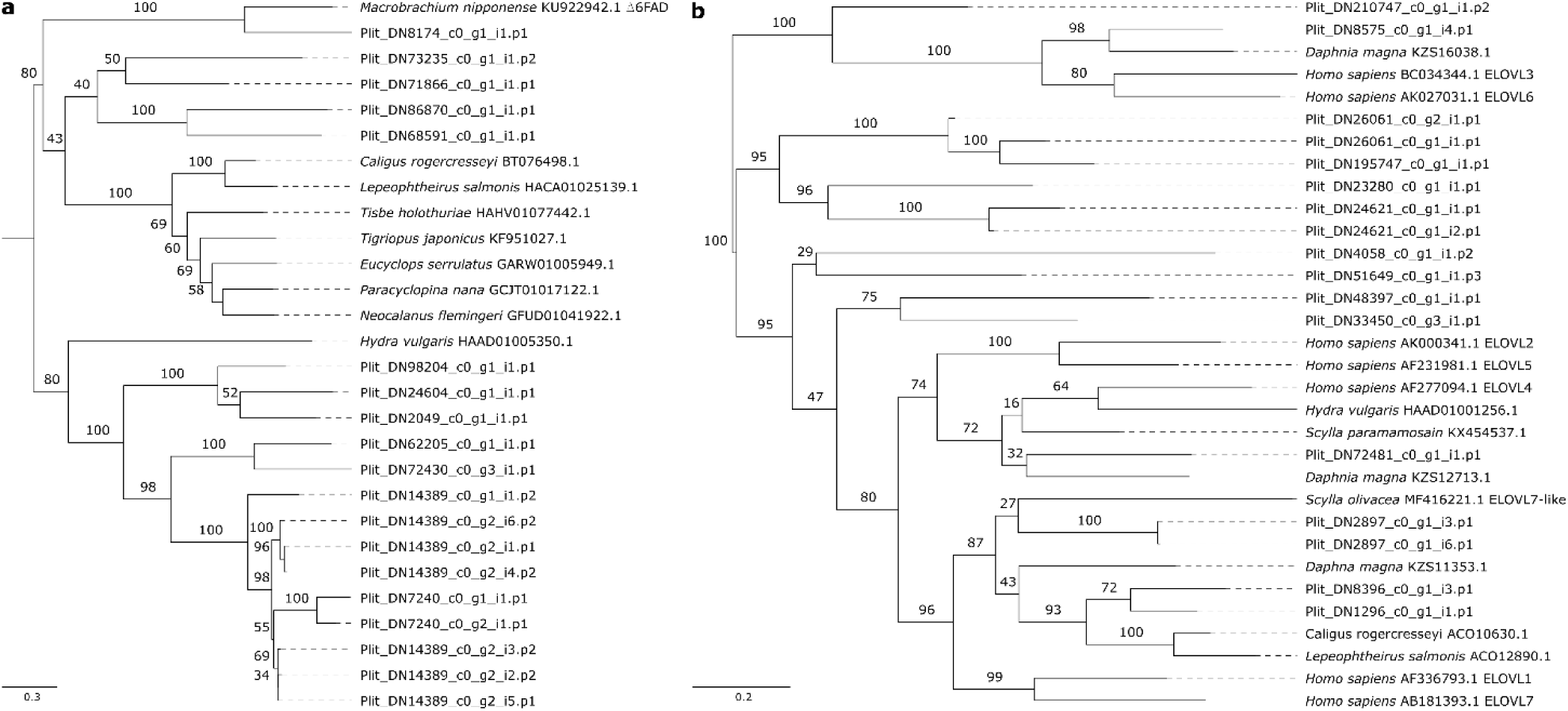
Maximum likelihood phylogenetic trees with midpoint root comparing trimmed nucleotide sequences of putative front-end desaturase (**a**) and elongase (**b**) transcripts of *P. littoralis* with sequences of other crustaceans, *Homo sapiens* and the hydrozoan *Hydra vulgaris*. Values above branches show bootstrap support after 100 RAxML iterations. Gene identification, when already functionally characterized, was added after each accession number.

## Discussion

Several earlier studies indicated that harpacticoid copepods have a strong capacity for endogenous LC-PUFA bioconversion (18,19,21). Meanwhile, our knowledge on the molecular pathways underlying this process is still limited (16). Considering a global SST increase and a decline in LC-PUFA production by primary consumers within this century (7), LC-PUFA bioconversion potentially plays a unique role for primary consumers to physiologically mitigate the negative effects of climate change. Prior to our experiment, the algae were lyophilized to prevent temperature-dependent growth. Previous research found that harpacticoids ingest lyophilized algae, albeit at a lower assimilation rate than live algae (58). We showed that when fed with the LC-PUFA-deficient *D. tertiolecta, P. littoralis* had the physiological plasticity to maintain LC-PUFA levels relatively similar to copepods fed with *Nitzschia* sp. The high abundance of ALA in *D. tertiolecta*-fed *P. littoralis* indicates that the copepods efficiently ingested lyophilized *D. tertiolecta*. This likely counteracted the lack of LC-PUFA, since *P. littoralis* is able to use ALA as a precursor for desaturation and elongation towards EPA and DHA (16). *D. tertiolecta*-derived 16:4*ω*3 and 16:3*ω*3 and *Nitzschia* sp.-derived 16:1*ω*7 were not sustained in *P. littoralis* (Supplementary Tables). Either way the lyophilized algae were not as efficiently assimilated as previously detected, or these compounds were bioconverted to other FAs or metabolized to provide energy. In contrast to diet, an increased SST of 3°C reduced the absolute total FA content, suggesting an increased use of storage FAs as energy providers (9). Some FA compounds showed extensive variability between replicates, which can be explained by the low sample volume or the low number of replicates. While the absolute concentrations of DHA and several monounsaturated FAs decreased, the relative concentrations of LC-PUFAs remained unaltered. Homeoviscous adaptation as a possible explanation cannot be confirmed nor rejected, as we do not have any information on membrane FAs specifically (6). Decreased DHA concentrations can alternatively be linked to an increased stress response at the upper limits of *P. littoralis*’ thermal range. In a previous experiment, increased temperatures indeed stimulated ARA and EPA bioconversion to allow enhanced eicosanoid biosynthesis in *P. littoralis* (19). However, a comparison with the current results should be done with care since the previous experiment included an ecologically improbable SST increase of 10°C. These results thus clearly illustrate the importance of LC-PUFA bioconversion as a mechanism to cope with the direct and indirect effects of global warming. As such, fluctuations in temperature or food quality might be relatively less detrimental to *P. littoralis*, and its historical adaptation to a variable environment could have given this species useful adaptations to persist under future environmental changes.

To investigate the molecular mechanisms behind the FA metabolism of *P. littoralis*, we performed a *de novo* assembly of its transcriptome. The high N50 metric (1,360) and high number of complete arthropod BUSCO genes (97.9 %) compared to other copepod transcriptomes lead us to state that the transcriptome presented here is of high quality and can be confidently used for future studies (59). In our differential gene expression analysis, seven and 29 transcripts were found to be DE in the dietary and temperature contrasts, respectively. These low numbers can be attributed to the employed filtering threshold, or to the low number of replicates used, which reduced the power to detect significantly DE genes. Low numbers of DE genes were found in other transcriptomic studies on copepods as well (59). A gene set enrichment analysis identified GO terms mainly related to cytoskeleton organisation and phospholipid biosynthesis. Possible relations between temperature or dietary LC-PUFA availability and the cytoskeleton have been identified in mammals (60,61), yet further investigations on invertebrates are needed. GO terms related to phospholipid biosynthesis were all attributed to one transcript which was downregulated when *P. littoralis* was fed *D. tertiolecta* and upregulated at 22°C, respectively. A temperature-driven reduction of membrane-bound phospholipid biosynthesis seems plausible (6) and is in line with both our own and previous findings (19) at the FA level. It may be interpreted as increased mobilization of FAs for energy provision. The previous study however did not find membrane FA depletion when *P. littoralis* was fed the LC-PUFA deficient diet *D. tertiolecta* (19), thereby contradicting our findings at the transcriptional level. As we sequenced bulk RNA from a pool of specimens, we were unable to determine whether some of the DE genes are tissue-specific regulated or not. Importantly, we identified a high number of transcripts putatively encoding for front-end desaturases and elongases. While we are aware that a *de novo* assembly may artificially introduce an inflated number of contigs (62), most sequences were sufficiently distinct to confidently state that they belong to different paralogous genes. It is indeed known that gene duplication is an important driver of the diversity of desaturase and elongase genes (63,64). We would therefore like to advance the hypothesis that an elevated front-end desaturase and elongase gene duplication frequency in harpacticoid copepods could be the key element for their high LC-PUFA bioconversion capacity. However, more exhaustive phylogenetic analyses are necessary to test this hypothesis. Our current analyses already show that putative front-end desaturases from *P. littoralis* grouped into two distinct phylogenetic clades, with sequences clustering with either way other copepod desaturase sequences and a Δ6 desaturase sequence from *M. nipponense*, or with a desaturase sequence from *H. vulgaris* (Fig. 4a). Corroborating the results of a previous study, copepod front-end desaturases grouped together in one monophyletic clade, however none of the *P. littoralis* sequences were found to cluster within this clade, but rather grouped separately as a sister clade. Including more functionally characterized front-end desaturases from other crustacean species might benefit our analyses, but are so far not available (16). In contrast, the maximum-likelihood tree of the elongase sequences corresponds more to earlier studies (27) (Fig. 4b). While ten *P. littoralis* sequences did not cluster with any of the additional sequences, we were able to assign seven *P. littoralis* sequences as well as seven other crustacean sequences to one of the three subclades Elovl3/Elvol6-like, Elovl4-like or Elovl1/Elovl7-like. Overall, our phylogenetic analyses of the front-end desaturase and elongase transcripts of *P. littoralis* provide a first glimpse at the high diversity of these genes and serve as a starting point to better comprehend the evolutionary history of LC-PUFA bioconversion within harpacticoid copepods. Interestingly, none of the identified transcripts encoding a front-end desaturase or elongase were found to be DE due to temperature or dietary LC-PUFA availability. This contradicts the stressor-driven LC-PUFA bioconversion evidenced at the FA level. The majority of those transcripts (31 out of 36) were classified as lowly expressed (<5 counts per million sample^−1^) and were therefore retained from the DE analysis. Additionally, the applied correction for multiple testing might have resulted in potentially significantly DE transcripts to go undetected (54). A more gene-specific approach such as reverse transcriptase quantitative polymerase chain reaction (RT-qPCR) (65) might be better suited to analyse expression of LC-PUFA bioconversion genes following direct and indirect effects of global warming.

In conclusion, we combined two approaches – FA profiling and *de novo* transcriptome assembly – to expand the current knowledge on LC-PUFA bioconversion in harpacticoids. This study shows that LC-PUFA levels in *P. littoralis* remain high even on a LC-PUFA-deficient diet, yet transcripts putatively encoding for front-end desaturases and elongases were not found to be upregulated. The molecular pathways underlying this mechanism are thus more complex than previously assumed (also demonstrated by the recent discovery of *ω*-end desaturases in multiple aquatic invertebrates (28)) and might not happen at the gene expression level. The *de novo* transcriptome of a non-model harpacticoid copepod presented here lays the foundation for more targeted ecophysiological research to investigate the molecular basis of adaptions to cope with the effects of global change.

## Acknowledgements

We thank Bruno Vlaeminck for his help with the fatty acid analysis. This work was supported by the Special Research Fund of Ghent University through a starting grant (BOF16/STA/028) and a GOA grant (01GA2617) and carried out with infrastructure provided by EMBRC Belgium (FWO GOH3817N). The first author is supported by a PhD grant fundamental research of the Research Foundation Flanders – FWO (11E2320N).

