## Supplementary Methods for "Fatty acid bioconversion in harpacticoid copepods in a changing environment: a transcriptomic approach"

**RNA extraction**

At the end of the experiment, after mortality assessment and cleaning in filtered and autoclaved natural seawater (FNSW) to remove food particles, *P. littoralis* individuals from each replicate were pooled and transferred to a 1.5 ml Eppendorf tube. Copepods were concentrated by allowing them to settle and leftover FNSW in the Eppendorf was removed. Immediately thereafter (within minutes after removal from the Lovibond incubator), samples were flash-frozen in liquid nitrogen prior to storage at -80°C. Total RNA was extracted using the RNeasy Plus Micro Kit (QIAGEN), with slight modifications to the manufacturer’s protocol. Samples were disrupted and simultaneously homogenized in 350 µl Buffer RTL using one stainless steel bead (5 mm) per sample and a TissueLyser II (Qiagen) for two times 45 s at 30 Hz, with the samples placed on ice for one minute after each disruption step. The sample was centrifuged for 3 min at 8,000 g, after which the lysate was transferred to a new 1.5 ml microcentrifuge by pipetting. 350 µl 70 % ethanol was added and mixed with lysate, and the mixture (700 µl) was immediately transferred to an RNeasy MinElute spin column placed in a 2 ml collection tube. The spin column was centrifuged for 15 s at 8,000 g, 350 µl Buffer RW1 was added, and the spin column was centrifuged again for 15 s at 8,000 g. A mixture of DNase I solution (10 µl) and Buffer RDD (70 µl) was added to the spin column for 15 min. The subsequent extraction steps were similar to the ones in the manufacturer’s protocol. At the final step, RNase-free water (19 µl) was added to the spin column and the spin column was centrifuged for 1 min at 8,000 g, resulting in a total RNA solution with a volume of about 17 µl. After quality and quantity assessment using a NanoDrop 2000 spectrophotometer (Thermo Scientific) and a 2100 Bioanalyzer (Agilent Technologies), samples (15 µl) were diluted to ensure equal total RNA concentrations prior to cDNA library construction. Final total RNA concentration per sample (17 µl) was 79.93 ± 6.68 ng µl^-1^.
