## Supplementary Tables for "Fatty acid bioconversion in harpacticoid copepods in a changing environment: a transcriptomic approach"

**Table S1** Mean relative FA concentration (% ± s.d.; n = 4) of *Nitzschia* sp. and *D. tertiolecta*

| Fatty acid | *Nitzschia* sp. | *D. tertiolecta* |
| --- | --- | --- |
| 14:0 | 9.09 ± 0.29 | 0.35 ± 0.03 |
| iso-15:0 | 0.19 ± 0.06 | 0.06 ± 0.04 |
| anteiso-15:0 | 0.10 ± 0.03 | 0.01 ± 0.01 |
| 15:0 | 0.54 ± 0.09 | n.d. |
| iso-16:0 | n.d. | 0.01 ± 0.01 |
| 16:0 | 9.07 ± 0.55 | 18.65 ± 0.78 |
| 16:1 | 3.46 ± 0.10 | 2.03 ± 0.14 |
| 16:1w7 | 10.14 ± 0.38 | 0.21 ± 0.07 |
| 16:2w4 | 6.20 ± 0.14 | n.d. |
| 16:2w6 | n.d. | 0.69 ± 0.05 |
| 16:2w7 | 1.53 ± 0.10 | n.d. |
| 16:3w3 | 0.07 ± 0.02 | 3.66 ± 0.34 |
| 16:3w4 | 3.59 ± 0.93 | 0.05 ± 0.02 |
| 16:3w6 | n.d. | 0.65 ± 0.11 |
| 16:4w1 | 7.14 ± 0.63 | 0.02 ± 0.01 |
| 16:4w3 | n.d. | 18.37 ± 0.81 |
| 18:0 | 0.91 ± 0.26 | 0.43 ± 0.10 |
| 18:1w7 | 8.95 ± 2.10 | n.d. |
| 18:1w9 | 0.14 ± 0.05 | 2.97 ± 0.46 |
| 18:2w6 | 0.65 ± 0.15 | 3.13 ± 0.37 |
| 18:3w3 | 0.05 ± 0.03 | 43.77 ± 1.29 |
| 18:3w6 | 1.82 ± 0.38 | 2.92 ± 0.22 |
| 18:4w3 | 1.26 ± 0.29 | 1.80 ± 0.06 |
| 20:4w6 | 0.41 ± 0.08 | 0.18 ± 0.02 |
| 20:5w3 | 26.85 ± 2.26 | n.d. |
| 22:0 | 0.07 ± 0.01 | 0.01 ± 0.01 |
| 22:5w3 | 0.13 ± 0.01 | 0.01 ± 0.00 |
| 22:6w3 | 5.68 ± 0.29 | n.d. |
| 24:0 | 1.98 ± 0.25 | 0.03 ± 0.00 |
| ∑SFA | 21.94 ± 0.91 | 19.54 ± 0.95 |
| ∑MUFA | 22.68 ± 2.44 | 5.22 ± 0.66 |
| ∑PUFA | 55.39 ± 3.34 | 75.24 ± 1.51 |

n.d. = not detected

**Table S2** Mean absolute FA concentration (ng copepod^-1^ ± s.d.; n = 3) of *P. littoralis* prior (field) and after ten days of incubation with *Nitzschia* sp. or *D. tertiolecta*

|  | Field |  | *Nitzschia* sp. | |  | *D. tertiolecta* | |  | Two-way ANOVA/Scheirer-Ray-Hare test | | |
| --- | --- | --- | --- | --- | --- | --- | --- | --- | --- | --- | --- |
|  |  |  | 19°C | 22°C |  | 19°C | 22°C |  | Diet | Temperature | Interaction |
| 14:0 | 8.94 ± 0.23 |  | 6.84 ± 3.65 | 3.61 ± 0.32 |  | 4.86 ± 0.25 | 4.19 ± 0.62 |  | - | - | - |
| iso-15:0 | 0.28 ± 0.04 |  | 1.43 ± 0.41 | 1.01 ± 0.11 |  | 1.67 ± 0.40 | 1.41 ± 0.26 |  | - | - | - |
| anteiso-15:0* | 0.15 ± 0.01 |  | 0.36 ± 0.26 | 0.15 ± 0.03 |  | 0.32 ± 0.16 | 0.19 ± 0.02 |  | - | p < 0.01 | - |
| 15:0 | 2.58 ± 0.20 |  | 2.49 ± 0.99 | 1.38 ± 0.11 |  | 2.19 ± 0.30 | 1.80 ± 0.22 |  | - | p < 0.05 | - |
| iso-16:0 | 0.34 ± 0.04 |  | 0.64 ± 0.48 | 0.24 ± 0.02 |  | 0.38 ± 0.07 | 0.27 ± 0.04 |  | - | - | - |
| 16:0 | 42.63 ± 0.90 |  | 38.88 ± 9.96 | 26.21 ± 1.44 |  | 34.82 ± 2.82 | 31.22 ± 3.29 |  | - | p < 0.05 | - |
| iso-17:0 | 0.38 ± 0.45 |  | 2.47 ± 1.91 | 0.85 ± 0.19 |  | 1.35 ± 0.08 | 1.23 ± 0.24 |  | - | - | - |
| 16:1w7* | 30.66 ± 1.31 |  | 19.82 ± 0.68 | 16.07 ± 0.89 |  | 19.91 ± 4.29 | 17.60 ± 1.60 |  | - | - | - |
| 16:1 | 0.19 ± 0.02 |  | 0.43 ± 0.10 | 0.19 ± 0.02 |  | 0.34 ± 0.24 | 0.31 ± 0.05 |  | - | - | - |
| 17:0 | 1.53 ± 0.12 |  | 1.83 ± 0.33 | 1.39 ± 0.05 |  | 1.60 ± 0.19 | 1.61 ± 0.14 |  | - | - | - |
| 16:2w4 | 2.36 ± 0.13 |  | 1.96 ± 0.86 | 1.36 ± 0.09 |  | 1.45 ± 0.34 | 1.76 ± 0.44 |  | - | - | - |
| 17:1 | 2.36 ± 0.13 |  | 1.96 ± 0.86 | 1.36 ± 0.09 |  | 1.45 ± 0.34 | 1.76 ± 0.44 |  | - | - | - |
| 18:0 | 7.29 ± 0.15 |  | 9.68 ± 4.94 | 6.00 ± 0.30 |  | 7.48 ± 0.85 | 7.00 ± 0.87 |  | - | - | - |
| 16:3w4 | 1.77 ± 0.15 |  | 1.59 ± 0.11 | 1.82 ± 0.19 |  | 1.67 ± 0.62 | 1.87 ± 0.19 |  | - | - | - |
| 18:1w7 | 9.37 ± 0.74 |  | 14.02 ± 0.79 | 11.23 ± 0.90 |  | 13.98 ± 1.42 | 12.08 ± 1.63 |  | - | p < 0.01 | - |
| 18:1w9 | 2.80 ± 0.09 |  | 4.89 ± 2.67 | 1.33 ± 0.13 |  | 2.36 ± 0.17 | 3.43 ± 2.21 |  | - | - | p < 0.05 |
| 16:4w1* | 1.65 ± 0.08 |  | 1.50 ± 0.34 | 1.22 ± 0.10 |  | 1.32 ± 0.50 | 1.22 ± 0.08 |  | - | - | - |
| 18:2w6 | 1.91 ± 0.11 |  | 1.11 ± 0.14 | 0.85 ± 0.04 |  | 1.21 ± 0.32 | 1.28 ± 0.22 |  | - | - | - |
| 18:3w6 | 1.25 ± 0.03 |  | 0.72 ± 0.10 | 0.59 ± 0.04 |  | 0.83 ± 0.30 | 0.79 ± 0.02 |  | - | - | - |
| 20:0 | 0.22 ± 0.03 |  | 0.43 ± 0.41 | 0.15 ± 0.02 |  | 0.30 ± 0.10 | 0.20 ± 0.07 |  | - | - | - |
| 18:3w3 | 0.83 ± 0.10 |  | 1.20 ± 0.72 | 0.50 ± 0.04 |  | 3.37 ± 0.85 | 3.11 ± 0.50 |  | p < 0.001 | - | - |
| 18:4w3 | 2.76 ± 0.07 |  | 1.46 ± 0.23 | 1.19 ± 0.08 |  | 1.49 ± 0.45 | 1.50 ± 0.14 |  | - | - | - |
| 22:0* | 1.74 ± 0.49 |  | 1.43 ± 0.06 | 1.19 ± 0.04 |  | 1.45 ± 0.18 | 1.30 ± 0.10 |  | - | - | - |
| 20:4w6 | 3.85 ± 0.24 |  | 2.89 ± 0.68 | 2.64 ± 0.23 |  | 2.76 ± 0.58 | 2.58 ± 0.15 |  | - | - | - |
| 20:5w3* | 44.78 ± 1.28 |  | 27.99 ± 3.11 | 24.22 ± 1.36 |  | 27.06 ± 5.50 | 25.01 ± 2.51 |  | - | - | - |
| 24:0 | 2.56 ± 0.11 |  | 2.68 ± 0.16 | 2.21 ± 0.05 |  | 2.31 ± 0.24 | 2.19 ± 0.16 |  | - | p < 0.05 | - |
| 22:5w3 | 3.24 ± 0.14 |  | 2.24 ± 0.05 | 1.80 ± 0.09 |  | 2.23 ± 0.81 | 1.81 ± 0.29 |  | - | - | - |
| 22:6w3 | 18.88 ± 0.08 |  | 17.40 ± 2.64 | 15.26 ± 0.84 |  | 17.42 ± 1.99 | 15.22 ± 0.11 |  | - | p < 0.05 | - |
| ∑SFA | 68.64 ± 2.33 |  | 69.14 ± 23.39 | 44.37 ± 2.26 |  | 58.73 ± 1.83 | 52.60 ± 5.88 |  | - | - | - |
| ∑MUFA | 45.38 ± 0.84 |  | 41.12 ± 4.17 | 30.17 ± 0.15 |  | 38.04 ± 5.89 | 35.18 ± 4.07 |  | - | p < 0.05 | - |
| ∑PUFA | 83.27 ± 2.13 |  | 60.07 ± 7.25 | 51.47 ± 2.34 |  | 60.82 ± 12.01 | 56.16 ± 3.78 |  | - | - | - |
| DHA/EPA* | 0.42 ± 0.01 |  | 0.62 ± 0.03 | 0.63 ± 0.04 |  | 0.65 ± 0.07 | 0.61 ± 0.07 |  | - | - | - |
| ∑FA | 197.29 ± 4.50 |  | 170.33 ± 17.76 | 126.02 ± 4.63 |  | 157.59 ± 19.55 | 143.94 ± 13.05 |  | - | p < 0.05 | - |

Differences were tested using type II two-way ANOVA, except for datasets without normal distribution (*), for which the Scheirer-Ray-Hare test was used.

- : not significant

**Table S3** Mean relative FA concentration (% ± s.d.; n = 3) of *P. littoralis* prior (field) and after ten days of incubation with *Nitzschia* sp. or *D. tertiolecta*

|  | Field |  | *Nitzschia* sp. | |  | *D. tertiolecta* | |  | Two-way ANOVA/Scheirer-Ray-Hare test | | |
| --- | --- | --- | --- | --- | --- | --- | --- | --- | --- | --- | --- |
|  |  |  | 19°C | 22°C |  | 19°C | 22°C |  | Diet | Temperature | Interaction |
| 14:0 | 4.53 ± 0.06 |  | 3.91 ± 1.71 | 2.86 ± 0.16 |  | 3.12 ± 0.42 | 2.9 ± 0.19 |  | - | - | - |
| iso-15:0 | 0.14 ± 0.02 |  | 0.83 ± 0.17 | 0.80 ± 0.08 |  | 1.08 ± 0.36 | 0.98 ± 0.12 |  | - | - | - |
| anteiso-15:0* | 0.08 ± 0.00 |  | 0.20 ± 0.13 | 0.12 ± 0.02 |  | 0.22 ± 0.14 | 0.13 ± 0.00 |  | - | - | - |
| 15:0 | 1.31 ± 0.08 |  | 1.43 ± 0.43 | 1.10 ± 0.12 |  | 1.42 ± 0.35 | 1.25 ± 0.04 |  | - | - | - |
| iso-16:0 | 0.17 ± 0.02 |  | 0.36 ± 0.24 | 0.19 ± 0.01 |  | 0.24 ± 0.08 | 0.19 ± 0.03 |  | - | - | - |
| 16:0* | 21.61 ± 0.11 |  | 22.63 ± 3.66 | 20.79 ± 0.39 |  | 22.18 ± 1.06 | 21.67 ± 0.61 |  | - | - | - |
| iso-17:0 | 0.19 ± 0.23 |  | 1.39 ± 0.96 | 0.67 ± 0.12 |  | 0.87 ± 0.16 | 0.85 ± 0.13 |  | - | - | - |
| 16:1w7 | 15.54 ± 0.44 |  | 11.74 ± 1.50 | 12.75 ± 0.45 |  | 12.53 ± 1.24 | 12.23 ± 0.01 |  | - | - | - |
| 16:1 | 0.10 ± 0.01 |  | 0.26 ± 0.08 | 0.15 ± 0.02 |  | 0.21 ± 0.14 | 0.22 ± 0.06 |  | - | - | - |
| 17:0 | 0.78 ± 0.04 |  | 1.07 ± 0.09 | 1.10 ± 0.05 |  | 1.02 ± 0.07 | 1.12 ± 0.04 |  | - | - | - |
| 16:2w4 | 1.20 ± 0.05 |  | 1.16 ± 0.50 | 1.08 ± 0.07 |  | 0.91 ± 0.11 | 1.22 ± 0.28 |  | - | - | - |
| 17:1 | 1.20 ± 0.05 |  | 1.16 ± 0.50 | 1.08 ± 0.07 |  | 0.91 ± 0.11 | 1.22 ± 0.28 |  | - | - | - |
| 18:0 | 3.69 ± 0.01 |  | 5.55 ± 2.30 | 4.76 ± 0.16 |  | 4.85 ± 1.22 | 4.86 ± 0.23 |  | - | - | - |
| 16:3w4 | 0.90 ± 0.07 |  | 0.95 ± 0.15 | 1.45 ± 0.15 |  | 1.04 ± 0.28 | 1.30 ± 0.02 |  | - | p < 0.01 | - |
| 18:1w7 | 4.76 ± 0.49 |  | 8.31 ± 1.17 | 8.93 ± 0.97 |  | 8.9 ± 0.56 | 8.38 ± 0.60 |  | - | - | - |
| 18:1w9* | 1.42 ± 0.08 |  | 2.82 ± 1.48 | 1.05 ± 0.09 |  | 1.52 ± 0.29 | 2.37 ± 1.48 |  | - | - | p < 0.05 |
| 16:4w1 | 0.84 ± 0.03 |  | 0.90 ± 0.27 | 0.97 ± 0.05 |  | 0.82 ± 0.24 | 0.85 ± 0.04 |  | - | - | - |
| 18:2w6 | 0.97 ± 0.03 |  | 0.65 ± 0.01 | 0.67 ± 0.03 |  | 0.76 ± 0.12 | 0.89 ± 0.08 |  | p < 0.01 | - | - |
| 18:3w6 | 0.63 ± 0.01 |  | 0.43 ± 0.09 | 0.47 ± 0.04 |  | 0.51 ± 0.14 | 0.55 ± 0.04 |  | - | - | - |
| 20:0* | 0.11 ± 0.01 |  | 0.24 ± 0.21 | 0.12 ± 0.01 |  | 0.20 ± 0.10 | 0.14 ± 0.04 |  | - | - | - |
| 18:3w3 | 0.42 ± 0.05 |  | 0.68 ± 0.34 | 0.40 ± 0.04 |  | 2.12 ± 0.31 | 2.16 ± 0.23 |  | p < 0.001 | - | - |
| 18:4w3* | 1.40 ± 0.03 |  | 0.87 ± 0.19 | 0.95 ± 0.03 |  | 0.93 ± 0.19 | 1.04 ± 0.00 |  | - | - | - |
| 22:0 | 0.88 ± 0.24 |  | 0.85 ± 0.11 | 0.94 ± 0.02 |  | 0.92 ± 0.03 | 0.91 ± 0.06 |  | - | - | - |
| 20:4w6* | 1.95 ± 0.08 |  | 1.73 ± 0.51 | 2.09 ± 0.14 |  | 1.74 ± 0.16 | 1.80 ± 0.22 |  | - | - | - |
| 20:5w3 | 22.69 ± 0.14 |  | 16.62 ± 3.05 | 19.22 ± 0.50 |  | 17.05 ± 1.57 | 17.37 ± 0.23 |  | - | - | - |
| 24:0* | 1.30 ± 0.05 |  | 1.58 ± 0.10 | 1.75 ± 0.03 |  | 1.47 ± 0.06 | 1.52 ± 0.07 |  | p < 0.05 | - | - |
| 22:5w3* | 1.64 ± 0.03 |  | 1.33 ± 0.13 | 1.43 ± 0.06 |  | 1.39 ± 0.37 | 1.25 ± 0.12 |  | - | - | - |
| 22:6w3 | 9.57 ± 0.18 |  | 10.36 ± 2.30 | 12.11 ± 0.47 |  | 11.07 ± 0.17 | 10.64 ± 1.03 |  | - | - | - |
| ∑SFA* | 34.79 ± 0.58 |  | 40.05 ± 9.65 | 35.20 ± 0.59 |  | 37.58 ± 3.83 | 36.51 ± 1.36 |  | - | - | - |
| ∑MUFA | 23.01 ± 0.55 |  | 24.29 ± 3.11 | 23.96 ± 0.79 |  | 24.08 ± 0.84 | 24.41 ± 1.23 |  | - | - | - |
| ∑PUFA | 42.21 ± 0.16 |  | 35.67 ± 6.64 | 40.84 ± 0.50 |  | 38.34 ± 3.15 | 39.07 ± 0.95 |  | - | - | - |
| DHA/EPA* | 0.42 ± 0.01 |  | 0.62 ± 0.03 | 0.63 ± 0.04 |  | 0.65 ± 0.07 | 0.61 ± 0.07 |  | - | - | - |

Differences were tested using type II two-way ANOVA, except for datasets without normal distribution (*), for which the Scheirer-Ray-Hare test was used.

- : not significant

**Table S4** Tested contrasts and corresponding significantly differentially expressed transcripts

| Contrast | logFC | logCPM | F-statistic | p-value | FDR |
| --- | --- | --- | --- | --- | --- |
| Differentially expressed transcript |  |  |  |  |  |
| Diet |  |  |  |  |  |
| Plit_DN4535_c0_g1_i10 | -11.46 | 3.70 | 140.42 | 4.95E-07 | 0.01 |
| Plit_DN515_c0_g1_i2 | 22.86 | 2.67 | 384.29 | 2.38E-06 | 0.03 |
| Plit_DN3624_c0_g1_i4 | -10.94 | 2.40 | 91.79 | 3.27E-06 | 0.03 |
| Plit_DN5709_c0_g1_i3 | -11.60 | 2.63 | 72.75 | 8.94E-06 | 0.05 |
| Plit_DN1805_c0_g1_i18 | -10.39 | 2.89 | 69.50 | 1.09E-05 | 0.05 |
| Plit_DN1331_c0_g1_i7 | 3.23 | 3.47 | 53.37 | 1.16E-05 | 0.05 |
| Plit_DN2138_c0_g1_i4 | 10.63 | 1.95 | 77.39 | 1.42E-05 | 0.05 |
| Temperature |  |  |  |  |  |
| Plit_DN4535_c0_g1_i10 | 11.70 | 3.70 | 153.48 | 3.32E-07 | 0.01 |
| Plit_DN3007_c0_g1_i1 | -3.79 | 3.41 | 83.75 | 1.22E-06 | 0.01 |
| Plit_DN12745_c0_g1_i1 | -3.21 | 3.09 | 68.79 | 3.32E-06 | 0.02 |
| Plit_DN1805_c0_g1_i18 | 11.78 | 2.89 | 91.11 | 3.38E-06 | 0.02 |
| Plit_DN4651_c0_g1_i7 | -2.85 | 5.32 | 66.60 | 3.90E-06 | 0.02 |
| Plit_DN3624_c0_g1_i4 | -10.05 | 2.40 | 84.37 | 4.72E-06 | 0.02 |
| Plit_DN273_c0_g1_i2 | -1.59 | 5.41 | 63.41 | 4.98E-06 | 0.02 |
| Plit_DN6288_c0_g1_i1 | -6.03 | 7.12 | 61.63 | 5.74E-06 | 0.02 |
| Plit_DN63_c0_g1_i8 | 1.08 | 5.84 | 58.64 | 7.32E-06 | 0.02 |
| Plit_DN182_c0_g1_i2 | -3.19 | 7.07 | 55.79 | 9.34E-06 | 0.02 |
| Plit_DN2425_c0_g1_i12 | -2.47 | 5.65 | 53.50 | 1.14E-05 | 0.03 |
| Plit_DN1505_c0_g1_i5 | -2.37 | 6.95 | 50.93 | 1.45E-05 | 0.03 |
| Plit_DN6346_c0_g1_i4 | -3.57 | 7.85 | 49.69 | 1.63E-05 | 0.03 |
| Plit_DN1207_c0_g1_i4 | -2.03 | 3.84 | 48.85 | 1.77E-05 | 0.03 |
| Plit_DN182_c0_g1_i6 | -3.21 | 5.48 | 48.70 | 1.79E-05 | 0.03 |
| Plit_DN5040_c0_g1_i1 | -3.81 | 3.61 | 47.99 | 1.93E-05 | 0.03 |
| Plit_DN374_c0_g1_i1 | -7.88 | 3.06 | 47.61 | 2.00E-05 | 0.03 |
| Plit_DN1071_c0_g1_i2 | -3.06 | 4.96 | 47.56 | 2.01E-05 | 0.03 |
| Plit_DN23020_c0_g1_i5 | -1.54 | 7.27 | 47.23 | 2.08E-05 | 0.03 |
| Plit_DN12448_c0_g1_i2 | -2.25 | 4.50 | 47.11 | 2.10E-05 | 0.03 |
| Plit_DN5709_c0_g1_i3 | 9.94 | 2.63 | 58.92 | 2.18E-05 | 0.03 |
| Plit_DN2138_c0_g1_i4 | -10.53 | 1.95 | 68.48 | 2.28E-05 | 0.03 |
| Plit_DN6269_c0_g1_i4 | -1.84 | 4.13 | 44.73 | 2.68E-05 | 0.03 |
| Plit_DN123_c1_g1_i1 | 11.24 | 2.22 | 52.34 | 3.56E-05 | 0.04 |
| Plit_DN166_c0_g1_i2 | -2.73 | 6.28 | 41.84 | 3.66E-05 | 0.04 |
| Plit_DN1551_c0_g1_i3 | -1.39 | 4.28 | 41.04 | 4.00E-05 | 0.04 |
| Plit_DN651_c0_g1_i2 | -2.34 | 4.99 | 39.65 | 4.68E-05 | 0.04 |
| Plit_DN4966_c0_g1_i3 | -3.88 | 3.62 | 38.48 | 5.37E-05 | 0.05 |
| Plit_DN2223_c0_g1_i2 | -3.99 | 2.92 | 37.68 | 5.90E-05 | 0.05 |
| Diet (temperature fixed at 19°C) |  |  |  |  |  |
| Plit_DN4535_c0_g1_i10 | -11.64 | 3.70 | 183.77 | 1.46E-07 | 0.00 |
| Plit_DN1805_c0_g1_i18 | -10.33 | 2.89 | 89.00 | 3.74E-06 | 0.04 |
| Plit_DN5709_c0_g1_i3 | -10.73 | 2.63 | 81.66 | 5.44E-06 | 0.04 |
| Diet (temperature fixed at 22°C) |  |  |  |  |  |
| Plit_DN3624_c0_g1_i4 | -10.41 | 2.40 | 107.94 | 1.60E-06 | 0.04 |
| Plit_DN1598_c0_g1_i9 | 13.73 | 4.58 | 211.68 | 3.06E-06 | 0.04 |
| Temperature (diet fixed at *D. tertiolecta)* |  |  |  |  |  |
| Plit_DN4535_c0_g1_i10 | 11.76 | 3.70 | 183.83 | 1.46E-07 | 0.00 |
| Plit_DN3624_c0_g1_i4 | -9.97 | 2.40 | 99.79 | 2.27E-06 | 0.02 |
| Plit_DN1805_c0_g1_i18 | 11.02 | 2.89 | 96.89 | 2.58E-06 | 0.02 |
| Plit_DN1598_c0_g1_i9 | 13.73 | 4.58 | 158.20 | 7.74E-06 | 0.05 |
| Plit_DN5709_c0_g1_i3 | 9.91 | 2.63 | 71.05 | 9.89E-06 | 0.05 |
| Temperature by diet interaction |  |  |  |  |  |
| Plit_DN4535_c0_g1_i10 | 11.81 | 3.70 | 156.08 | 3.07E-07 | 0.01 |
| Plit_DN1598_c0_g1_i9 | 26.31 | 4.58 | 320.58 | 8.08E-07 | 0.01 |
| Plit_DN3624_c0_g1_i4 | -9.88 | 2.40 | 82.01 | 5.34E-06 | 0.04 |

Differential expression was tested using the edgeR gene-wise negative binomial generalized linear model. LogFC: log fold change; logCPM: log counts per million; FDR: false discovery rate.

**Table S5** Gene Ontology (GO) terms enriched under the diet and temperature treatments

| Domain | GO term | Fisher p-value |
| --- | --- | --- |
| GO.ID |  |  |
| Diet | | |
| Biological process |  |  |
| GO:0032053 | ciliary basal body organization | 0.0017 |
| GO:0010457 | centriole-centriole cohesion | 0.0022 |
| GO:1903566 | positive regulation of protein localization to cilium | 0.0026 |
| GO:0031053 | primary miRNA processing | 0.0050 |
| GO:0006654 | phosphatidic acid biosynthetic process | 0.0054 |
| GO:0030951 | establishment or maintenance of microtubule cytoskeleton polarity | 0.0066 |
| GO:0016024 | CDP-diacylglycerol biosynthetic process | 0.0072 |
| GO:0097711 | ciliary basal body-plasma membrane docking | 0.0075 |
| GO:0045724 | positive regulation of cilium assembly | 0.0076 |
| GO:0010669 | epithelial structure maintenance | 0.0096 |
| Molecular function |  |  |
| GO:0015232 | heme transporter activity | 0.0012 |
| GO:0030729 | acetoacetate-CoA ligase activity | 0.0012 |
| GO:0102420 | sw1-glycerol-3-phosphate C16:0-DCA-CoA acyl transferase activity | 0.0022 |
| GO:0004366 | glycerol-3-phosphate O-acyltransferase activity | 0.0023 |
| GO:0070878 | primary miRNA binding | 0.0026 |
| GO:0003841 | 1-acylglycerol-3-phosphate O-acyltransferase activity | 0.0041 |
| GO:0051010 | microtubule plus-end binding | 0.0049 |
| GO:0019894 | kinesin binding | 0.0085 |
| Cellular component |  |  |
| GO:0070877 | microprocessor complex | 0.0015 |
| GO:0035253 | ciliary rootlet | 0.0027 |
| GO:0005813 | centrosome | 0.0048 |
| GO:0001917 | photoreceptor inner segment | 0.0094 |
| Temperature | | |
| Biological process |  |  |
| GO:1903033 | positive regulation of microtubule plus-end binding | 0.0010 |
| GO:1904825 | protein localization to microtubule plus-end | 0.0012 |
| GO:0002149 | hypochlorous acid biosynthetic process | 0.0021 |
| GO:1990268 | response to gold nanoparticle | 0.0021 |
| GO:0034374 | low-density lipoprotein particle remodeling | 0.0025 |
| GO:0032053 | ciliary basal body organization | 0.0029 |
| GO:1905907 | negative regulation of amyloid fibril formation | 0.0030 |
| GO:0010457 | centriole-centriole cohesion | 0.0037 |
| GO:0001878 | response to yeast | 0.0041 |
| GO:1903566 | positive regulation of protein localization to cilium | 0.0043 |
| GO:0044130 | negative regulation of growth of symbiont in host | 0.0070 |
| GO:0045737 | positive regulation of cycliwdependent protein serine/threonine kinase activity | 0.0076 |
| GO:0031053 | primary miRNA processing | 0.0084 |
| GO:0002679 | respiratory burst involved in defense response | 0.0088 |
| GO:0006654 | phosphatidic acid biosynthetic process | 0.0091 |
| Molecular function |  |  |
| GO:0005509 | calcium ion binding | 0.0002 |
| GO:0015232 | heme transporter activity | 0.0027 |
| GO:0030021 | extracellular matrix structural constituent conferring compression resistance | 0.0029 |
| GO:0030729 | acetoacetate-CoA ligase activity | 0.0029 |
| GO:0008022 | protein C-terminus binding | 0.0034 |
| GO:0004035 | alkaline phosphatase activity | 0.0042 |
| GO:0102420 | sw1-glycerol-3-phosphate C16:0-DCA-CoA acyl transferase activity | 0.0051 |
| GO:0004366 | glycerol-3-phosphate O-acyltransferase activity | 0.0053 |
| GO:0070878 | primary miRNA binding | 0.0060 |
| GO:0003841 | 1-acylglycerol-3-phosphate O-acyltransferase activity | 0.0096 |
| Cellular component |  |  |
| GO:0070877 | microprocessor complex | 0.0025 |
| GO:0065010 | extracellular membrane-bounded organelle | 0.0040 |
| GO:0035253 | ciliary rootlet | 0.0045 |
| GO:0035371 | microtubule plus-end | 0.0071 |
| GO:0005796 | Golgi lumen | 0.0096 |
